# PecanPy: a fast, efficient, and parallelized Python implementation of *node2vec*

**DOI:** 10.1101/2020.07.23.218487

**Authors:** Renming Liu, Arjun Krishnan

## Abstract

Learning low-dimensional representations (embeddings) of nodes in large graphs is key to applying machine learning on massive biological networks. *Node2vec* is the most widely used method for node embedding. However, its original Python and C++ implementations scale poorly with network density, failing for dense biological networks with hundreds of millions of edges. We have developed PecanPy, a new Python implementation of *node2vec* that uses cache-optimized compact graph data structures and precomputing/parallelization to result in fast, high-quality node embeddings for biological networks of all sizes and densities. PecanPy software and documentation are available at https://github.com/krishnanlab/pecanpy.

## Background

Large-scale molecular networks are powerful models that capture interactions between biomolecules (genes, proteins, metabolites) on a genome scale (McGillivray *et al.*, 2018) and provide a basis for predicting novel associations between individual genes/proteins and various cellular functions, phenotypic traits, and complex diseases (Liu *et al.*, 2020; Sharan *et al.*, 2007). An area of research that has gained rapid adoption in network science across disciplines is learning low-dimensional numerical representations, or “embeddings”, of nodes in a network for easily leveraging machine-learning (ML) algorithms to analyze large networks (Goyal and Ferrara, 2018; Cai *et al.*, 2018; Hamilton *et al.*, 2018). Since each node’s embedding vector concisely captures its network connectivity, node embeddings can be conveniently used as feature vectors in any ML algorithm to learn/predict node properties or links (Hamilton et al., 2018). Most popular among the numerous node embedding methods proposed so far is *node2vec*, a random-walk based network embedding approach (Grover and Leskovec, 2016). Recent work has shown that *node2vec* has superior performance in node classification tasks on biological networks (Liu *et al.*, 2020; Yue *et al.*, 2020; Nelson *et al.*, 2019).

However, despite its popularity, the original *node2vec* software implementations (written in Python and C++) presents a significant bottleneck in seamlessly using *node2vec* on all current biological networks. First, due to inefficient memory usage and data structure, they do not scale to large and dense networks produced by integrating several data sources on a genome-scale (17–26k nodes and 3–300mi edges) (Greene *et al.*, 2015; Szklarczyk *et al.*, 2015). Next, the embarrassingly-parallel precomputations of calculating transition probabilities and generating random walks are not parallelized in the original software. Finally, the original implementations only support integer-type node identifiers (IDs), making it inconvenient to work with molecular networks typically available in databases where nodes may have non-integer IDs.

Recent work presented in preprints (Zhou *et al.*, 2018) and unpublished code repositories [1 2] have proposed improved implementations of *node2vec*. However, they either do not provide publicly-available software or do not present a full analysis of their implementation, including a benchmark that ensures the quality of the resulting embeddings. Here, we present PecanPy, an efficient Python implementation of *node2vec* that is parallelized, memory efficient, and accelerated using Numba with a cache-optimized data structure. We have extensively benchmarked our software using networks from the original study and multiple additional large biological networks to demonstrate both the computational performance and the quality of the node embeddings. In the rest of this paper, we first summarize the optimization and the performance of PecanPy and then go into the details of our implementation.

## Results

### Optimizing *node2vec* for Large Networks

Our goal was to optimize *node2vec* to make it work efficiently on a variety of networks that vary considerably in their sizes and densities, including large and dense graphs such as genome-scale gene interaction networks or microbial community sequence similarity networks. We chose to implement these improvements as a software in Python because it is currently the most widely-used high-level language in machine learning, making it convenient to use and to develop further as part of the community. Here, we present PecanPy, a new software for **p**arallelized, **e**ffi**c**ient, and **a**ccelerated ***n**ode2vec* in **Py**thon (**Figure 1**). PecanPy operates in three different modes – *PreComp*, *SparseOTF*, and *DenseOTF* – that are optimized for networks of different sizes and densities; *PreComp* for networks that are small and sparse, *SparseOTF* for networks that are large and sparse, and *DenseOTF* for dense networks. These modes appropriately take advantage of compact/dense graph data structures, precomputing transition probabilities, and computing 2nd-order transition probabilities during walk generation to achieve significant improvements in performance (**Table 1**). Next, we first showcase the performance of PecanPy before providing a detailed description of these optimizations.

**Figure 1.**
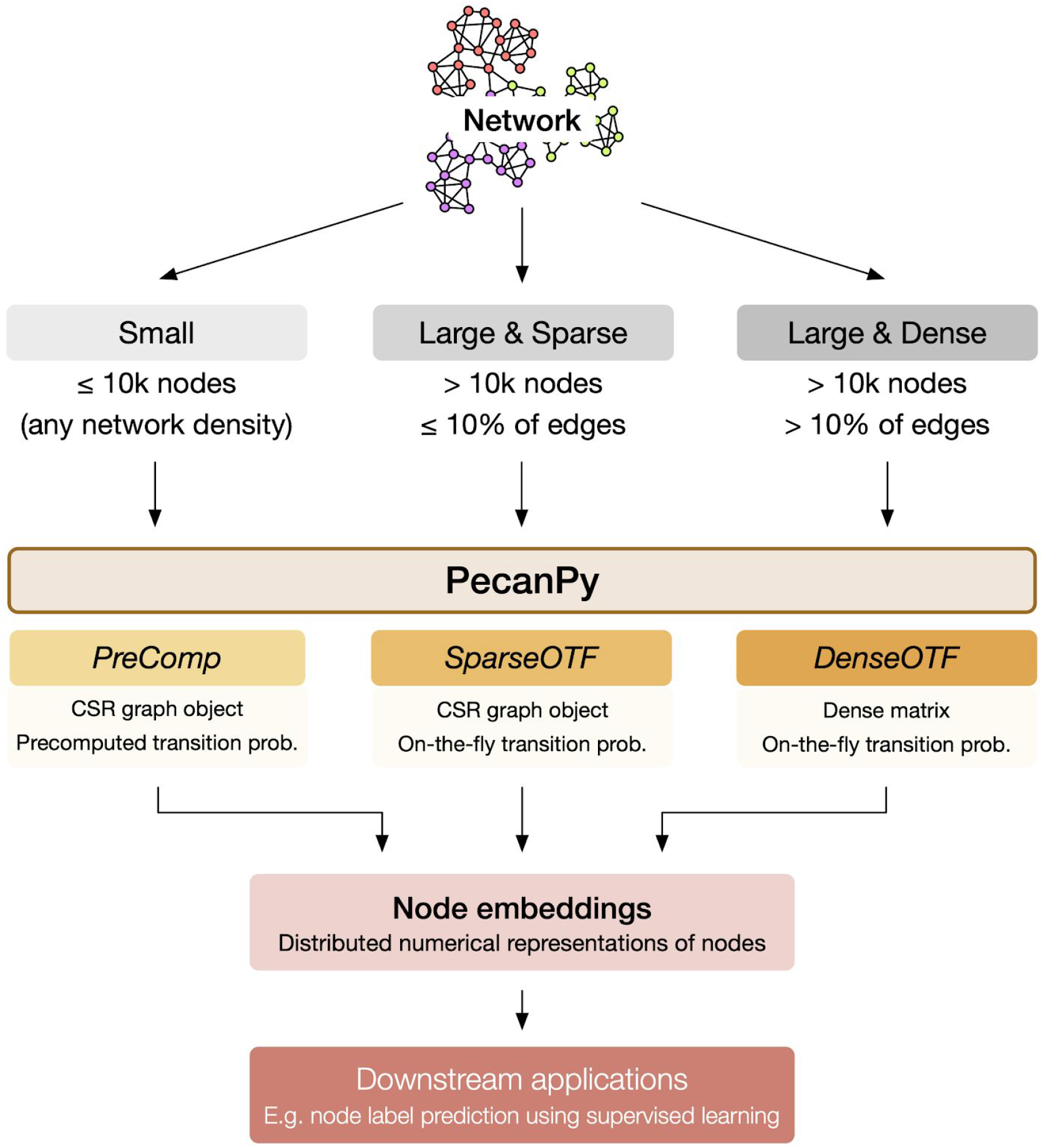
Overview *PecanPy* **for fast***node2vec*. PecanPy is a Python implementation of the *node2vec* algorithm. PecanPy can operate in three different modes – *PreComp*, *SparseOTF*, and *DenseOTF* – that are optimized for networks of different sizes and densities; *PreComp* for networks that are small (≤10,000 nodes; any density), *SparseOTF* for networks that are large and sparse (>10,000 nodes and ≤10% of possible edges, i.e. density ≤0.1), and *DenseOTF* for dense networks (>10,000 nodes and >10% of possible edges). These modes appropriately take advantage of compact sparse row (CSR) or dense matrix graph data structures, precomputing transition probabilities (prob.), and computing 2nd-order transition probabilities during walk generation to achieve significant improvements in performance.

**Table 1.** Summary of *node2vec* implementations.

| Implementation | Graph data structure | Precompute transition probabilities | Parallelized walk | Non-integer node ID |
| --- | --- | --- | --- | --- |
| Original Python | NetworkX | ✓ | x | x |
| Original C++ | SNAP | ✓ | ✓ | x |
| PreComp | CSR | ✓ | ✓ | ✓ |
| SparseOTF | CSR | x | ✓ | ✓ |
| DenseOTF | Dense Matrix | x | ✓ | ✓ |

### PecanPy Offers the Best *node2vec* Runtime and Memory Usage

We comprehensively benchmarked PecanPy and the original Python and C++ implementations of *node2vec* on a collection of eight networks (used throughout this study) including three networks from the original *node2vec* paper (Grover and Leskovec, 2016) and five large biological networks that together span a wide range of sizes (approx. 4k to 800k nodes and approx. 38k – 333mi edges) and densities (0.02% to 100%; **Table 2**; see *Methods* for more details).

**Table 2.**
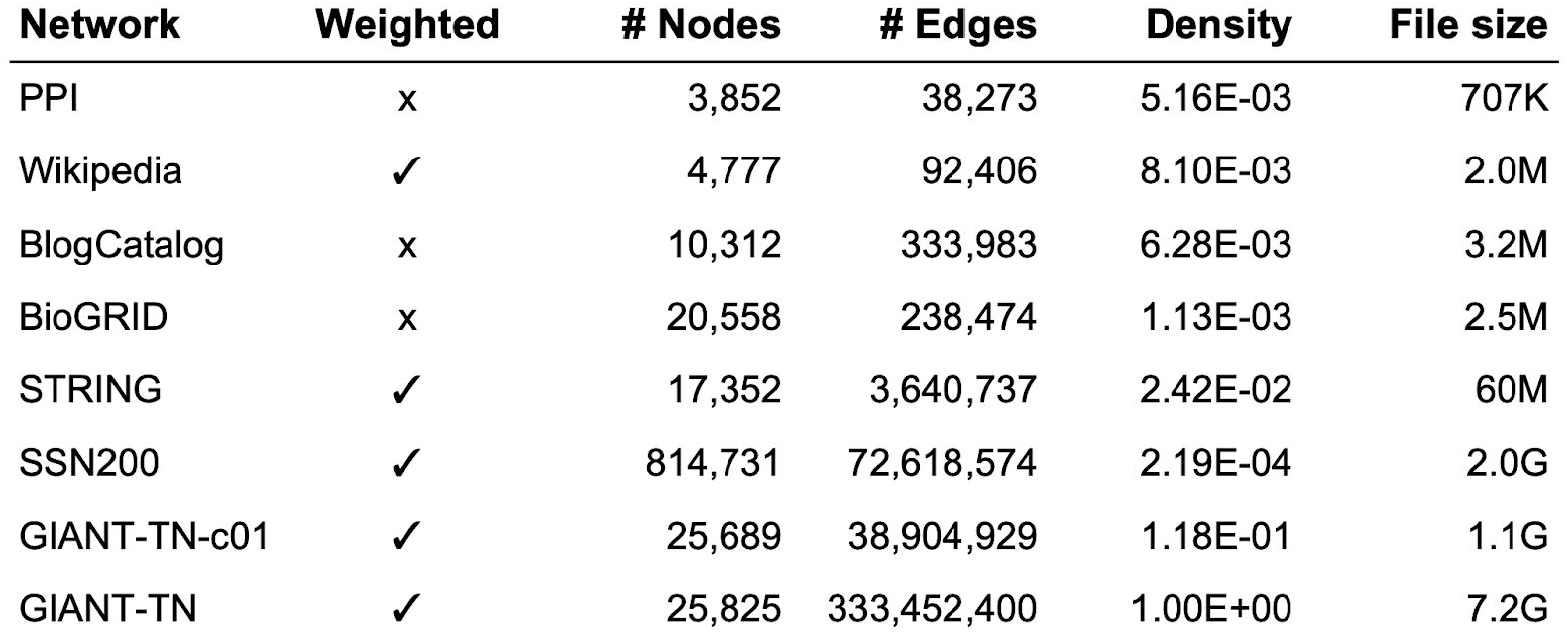
Properties of the diverse networks used in this study.

The performance of the two original (Original Python, Original C++) and the three new implementations (*PreComp*, *SparseOTF*, *DenseOTF*) in terms of runtime (**Fig. 2A**) and memory usage (maximum resident size; **Fig. 2B**) in a multi-core configuration are summarized in **Figure 2**. The full data is plotted in **Figure S1**. First, across the board, PecanPy (in one of its three modes) is substantially faster than the original implementations. In fact, there are three large networks (SSN200, GIANT-TN-c01, GIANT-TN) that run successfully only using PecanPy’s *OTF* implementations. The original software failed to run SSN200 because they do not support non-integer-type node IDs; this is supported in PecanPy. In terms of memory usage (**Fig. 2B** and **Fig. S1B**), one of the modes of the PecanPy reduces the maximum resident size by up to two orders of magnitude compared to the original implementations. These trends are magnified when all the implementations are run on a single core (**Fig. S2** and **S3**).

**Figure 2.**
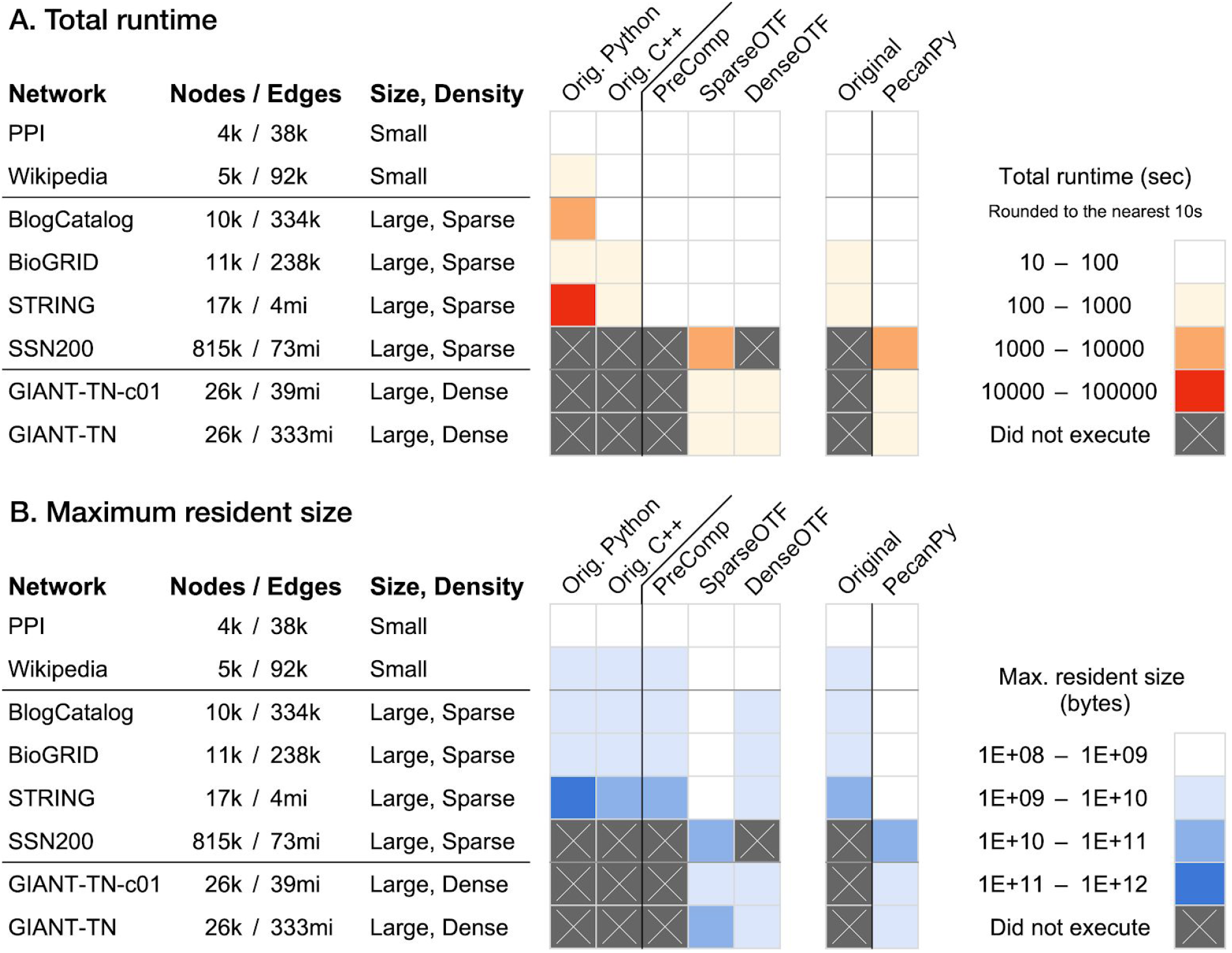
Summary of runtime and memory of PecanPy and the original implementations of *node2vec* using multiple cores. The eight networks of varying sizes and densities are along the rows. The software implementations are along the columns. The first heatmap (on the left) shows the performance of the original Python and C++ software along with the three modes of PecanPy (*PreComp*, *SparseOTF*, and *DenseOTF*). The adjacent 2-column heatmap (on the right) summarizes the performance of the original (best of Python and C++ versions) and PecanPy (best of *PreComp*, *SparseOTF*, and *DenseOTF*) implementations. Lighter colors correspond to lower runtime in panel A and lower memory usage in panel B. Crossed grey indicates that the particular implementation (column) failed to run for a particular network (row).

In the following sections, we outline how *node2vec* works, what the bottlenecks in the original software are, and how we addressed these bottlenecks.

### A Brief Description of *node2vec*

*Node2vec* (Grover and Leskovec, 2016) is a random-walk-based node embedding method (Hamilton *et al.*, 2018) that begins by generating a corpus of random walks on the input graph. In contrast to earlier approaches such as *DeepWalk* (Perozzi *et al.*, 2014) that perform 1st order random walk, *node2vec* performs 2nd order random walk where the transition probability from a current node to the next node also depends on the previous node that was visited. This 2nd order random walk strategy allows the flexibility of choosing between search strategies such as Breadth First Search and Depth First Search by setting the values of the return parameter and in-out parameter. These 2nd-order walks are then fed to the skip-gram model in the *word2vec* algorithm (Mikolov *et al.*, 2013) to compute the vector representation of each node.

The original *node2vec* software executes this method in four stages: loading, preprocessing, walking, and training. In the loading stage, graph edgelist files are read by the software and loaded into memory as graph objects. Then, during the preprocessing stage, all 2nd order transition probabilities are calculated and stored in memory for future reference. In the walking stage, walks are generated by randomly selecting the next node using the precomputed transition probabilities corresponding to current and previous states. Finally in the training stage, the generated walks are fed into the skip-gram model to generate the embeddings.

### Bottlenecks in the Original *node2vec* Implementation

To identify the bottlenecks in the original Python and C++ software, we profiled in detail the four stages of *node2vec* – loading, preprocessing, walking, and training – across the eight networks that vary widely in terms of size (number of nodes and edges), density, and edge weightedness (**Table 2**). Three of these networks – BlogCatalog, Wikipedia, and PPI – were used in the original *node2vec* paper (Grover and Leskovec, 2016).

For five out of the eight networks (**Fig. 3A-E**), among the four stages, training constituted dramatically different fractions of runtime between the Python and C++ implementations (first two stacked bars in each panel of **Figure 3**). Training only takes 1.2% (median) of the total runtime for the original Python implementation, in contrast to 95.1% for the original C++ implementation. Examining the raw training runtimes (**Fig. 4A**) shows that training the skip-gram using the *gensim* Python package is consistently an order of magnitude faster than the original C++ implementation. On the other hand, the walking and preprocessing stages in Python were considerably slower than in C++. Finally, both implementations failed to even load the three largest networks (**Fig. 3F-H** and **Fig. 4**). Naturally, all these bottlenecks are exacerbated when running these implementations on a single core (**Fig. S4** and **S5**).

**Figure 3.**
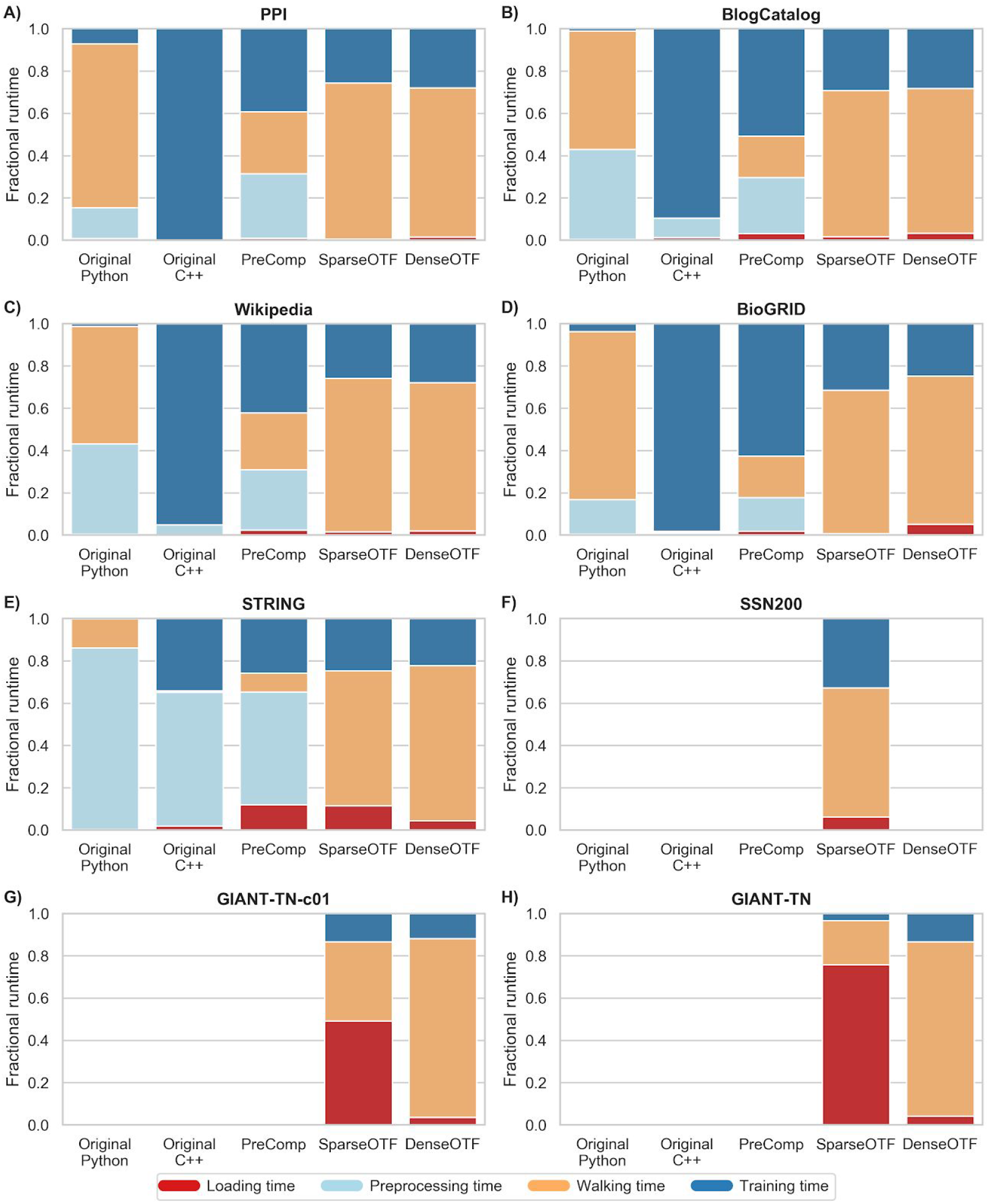
Fraction of runtime contributed by each stage of *node2vec* in different implementations using multiple cores. Each panel corresponds to a single network and each stacked bar within a panel corresponds to an individual *node2vec* implementation. The height of each segment within a bar represents the fraction of runtime contributed by each of the different stages of *node2vec*, tested in a multi-core configuration.

**Figure 4.**
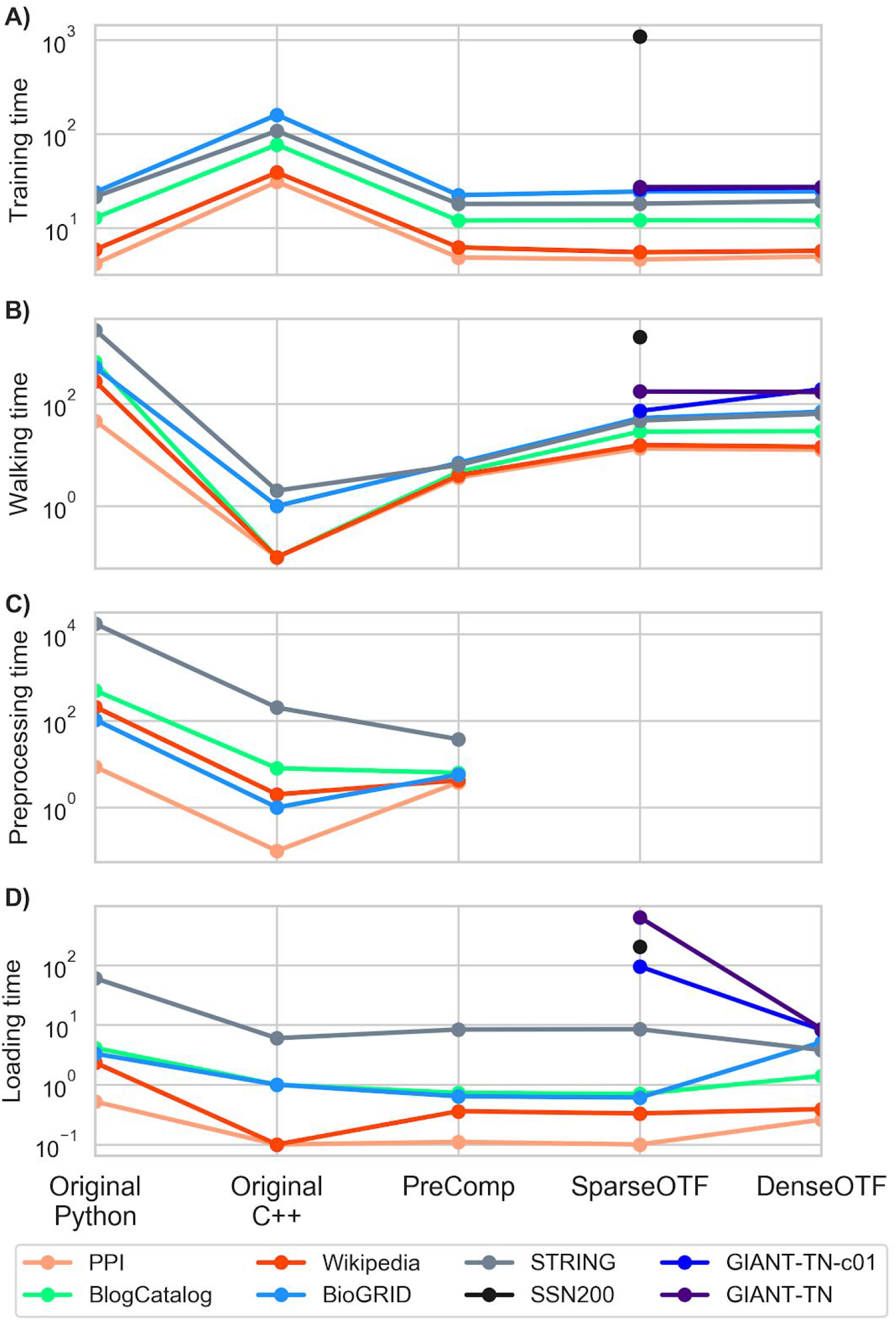
Raw runtimes of each stage of *node2vec* in different implementations using multiple cores. Each parallel plot corresponds to one of four stages of *node2vec*. Each line traces the raw runtime (points on parallel y-axes) of a specific network (color) across the different implementations (x-axis). In all plots, absence of a point for a particular network for any implementation indicates that the network failed to load.

First, as mentioned before, we picked Python due to its prevalence in machine learning. Next, based on the profiling results, we concentrated on optimizing the first three inefficient stages of *node2vec* in Python. Specifically, we concentrated on 1) implementing computationally- and memory-optimized graph data structures with efficient loading strategies; 2) providing an option to bypass the need to store all transition probabilities, leading to a significant reduction in memory usage; and 3) parallelizing the processes of transition probability computation and walk generation.

### Efficient Graph Data Structure to Improve Loading Networks

The first improvement we made was to implement a more efficient graph data structure for loading networks into the software. In the original Python implementation, *NetworkX*, which implicitly assumes that the input is a multigraph, was used to handle all operations on networks using a graph object in the form of nested dictionaries (dict-of-dict-of-dict; (Hagberg *et al.*, 2008)). The levels of these dictionaries correspond to node, neighbors of node, edge type, and edge weight. Thus, explicit declaration of edge types for weighted edges is required. However, *node2vec* only deals with homogeneous networks where only one type of edge is present in the network. In the original Python implementation, all edge types are set to “weight” by default. This extra piece of information requires 295 additional bytes of memory for every single edge stored in a *NetworkX* graph (empty dictionary = 240bytes, empty string = 49bytes, single character = 1byte). This requirement not only causes memory overheads, but also computation overheads by reading the dictionary for “edge type” that is irrelevant for homogeneous networks. To address these issues, in this work, we implemented a lite graph object as a network loader in the form of list-of-dict, which assumes the network has only one type of edge. As shown in **Figure 5** (first and second bars in each group), both the loading time and memory usage for list-of-dict were significantly reduced compared to that for *NetworkX*. Thus, the lite graph object efficiently loads networks with reduced memory usage and shorter load time compared to *NetworkX*.

**Figure 5.**
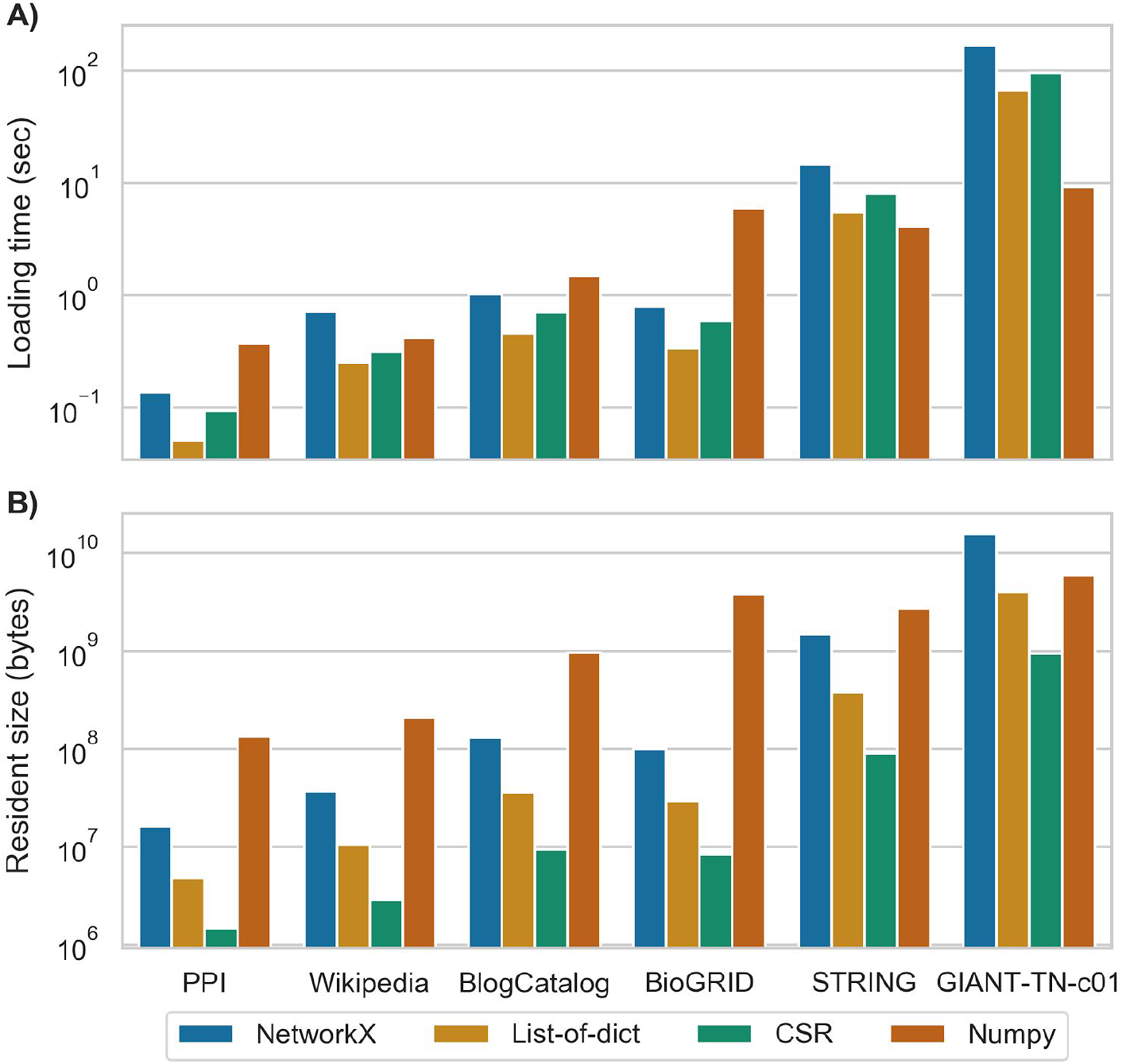
Effect of graph data structure on loading time and memory usage. Plot (A) shows the total time for loading networks, and plot (B) shows the total memory usage. Each plot contains groups of bars, each group corresponding to a network (among seven select networks), and each bar in the group corresponding to a specific network data structure.

### Cache Optimization to Further Optimize Graph Data Structure

Next, we optimized the graph data structure further for better cache utilization during computation. Despite being able to load faster with less memory usage using list-of-dict, operating on Python dictionaries is still suboptimal from the perspective of cache utilization. Specifically, the neighboring-edge data, which are often used together to compute transition probabilities, are not physically close to each other in memory. The design principle behind cache, however, is that units of memory that are physically nearby are likely to be used together. Consequently, every time a specific piece of data in memory is accessed, a chunk of physical memory that sits right next to the desired memory – called the cache line – is also copied from RAM to cache. Moreover, reading from cache could be up to 100 times faster than reading from RAM. To fully leverage the spatial locality of cache lines, inspired by a recent blog post (https://www.singlelunch.com/2019/08/01/700x-faster-node2vec-models-fastest-random-walks-on-a-graph/), we further converted the list-of-dict graph data structure to the compact sparse row (CSR) format, implemented using NumPy arrays (Walt *et al.*, 2011). In this way, neighboring-edge data is placed physically close together, thus improving cache utilization. *PreComp* uses CSR as their underlying graph data structure. These optimizations led to speedups in the preprocessing (**Fig. S2C**) and walk generation (**Fig. S2B**) steps for *PreComp* compared to that for the original Python implementation in the single-core setup. This includes up to an order-of-magnitude speedup for preprocessing by *PreComp* on both the BlogCatalog and Wikipedia networks.

Due to the compactness of the CSR representation of sparse matrices, memory usage can be further reduced compared to list-of-dict, as shown in **Figure 5B** (this bar in each group). However, for the same reason of compactness, dynamically constructing sparse matrices using CSR is extremely inefficient and expensive. Hence, we first use list-of-dict to load the network into memory and then convert the full network to CSR, on which the computation will be performed.

### Optimization for Dense Networks

The CSR representation described above is memory-efficient only if the network is relatively sparse because it stores non-zero entries of the adjacency matrix using both the indices and the weights of edges. For dense networks, explicit indexing of edges would cause memory overhead for the same reasons that redundancy of edge-type information (discussed before) would strain memory. To address this issue, for dense networks like GIANT-TN (25,825 nodes, fully connected and weighted; 333,452,400 edges), we implemented a dense NumPy matrix instead of CSR as the graph data structure.

CSR and dense matrix result in roughly similar walking times, as can be seen from comparing *SparseOTF* (that uses CSR) to *DenseOTF* (**Fig. 3B**). The benefit of using the dense matrix data structure over CSR instead comes from faster network loading and less memory usage. For dense networks, loading time takes up most of the runtime. For example, in the multi-core setup, loading the GIANT-TN network as CSR contributes to nearly 80% of the runtime (**Fig. 3H**). This burden is mostly due to the inefficiency in reading edgelist files as text line-by-line. To mitigate this issue of long loading time for dense networks, we implemented an option in our software that offers users the ability to first convert the edgelist file to dense matrix format and save as a binary Numpy npz file, which could then be loaded as a network in the future. As shown in **Figure 5A**, for sparse networks like PPI, loading networks as CSR is faster than loading as npz files, but as the density of network increases, loading networks as npz files becomes a better option. Similar arguments could be made for memory usage (**Fig. 5B**). For GIANT-TN (**Fig. S4D***SparseOTF* vs *DenseOTF*), loading as npz only took 9 seconds with 6GB of peak memory usage, resulting in a 70x speedup over CSR, which took more than 10 mins to load with 39GB of peak memory usage.

Then, the question is, when exactly is using a dense matrix more optimal than using CSR. A dense matrix requires 8 × *N*^2^ bytes of memory, while a CSR requires roughly 12 × *E* bytes of memory (using 32 bit unsigned integer as index and 64 bit floating point number as data), where *N* is the number nodes and *E* is the number of edges. Hence, in theory, for any network with density less than two-thirds, CSR should be used over the corresponding dense matrix. However, since CSR cannot be directly loaded but requires an intermediate conversion using list-of-dict, the peak memory usage is also affected by the list-of-dict graph data structure. Based on the observations from **Figure 5B**, where the fold difference in peak memory usage between CSR and Numpy matrix is much smaller for other sparse networks like BioGRID, we empirically set the balancing point for network density to be around 1/10.

### Memory Usage

Next, we focussed on reducing memory usage by computing 2^nd^ order transition probabilities on the fly. The original *node2vec* implementations precompute and store all the 2nd order transition probabilities in advance, which takes up at least 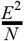 space in memory. One remedy for this issue with high memory usage is to calculate the 2^nd^ order transition probabilities On-The-Fly (*OTF*) during walk generation without saving them. Another reason for not precomputing 2nd order transition probabilities is that, as the network becomes larger and denser, it is very likely that most of the pre-calculated 2^nd^ order transition probabilities are never used in the generation of walks, causing not only the space, but also the invested computation time to be wasted. We combined the CSR and the dense matrix representation each with the *OTF* strategy into separate modes in our software called *SparseOTF* and *DenseOTF*, respectively. As an instance of improvement of the OTF strategy, the *DenseOTF* implementation was able to embed a dense network like the ~26k fully-connected weighted GIANT-TN network (>333 million edges) in just an hour with only 6GB of peak memory usage, using a single core (**Table S1**). Remarkably, this means that even for an extremely dense and large network like GIANT-TN, the software could be run on a personal computer configured with a reasonable amount of memory (e.g. 16GB).

Either of the original *node2vec* implementations failed to run even the sparsified version GIANT-TN-c01 (similar number of nodes and ~11% of the edges as GIANT-TN) on a supercomputer configured with 200GB memory. Moreover, the runtime of 1 hour for embedding the GIANT-TN network using *DenseOTF* with single-core configuration is even shorter than that for embedding a significantly smaller and sparser network STRING (67% of the nodes and 1% of edges as in GIANT-TN) using the original Python implementation with 28 cores, which took 5 hours to finish (**Table S1**).

For relatively small and sparse graphs where the amount of memory on a computer is sufficient to fit all 2nd-order transition probabilities, it could indeed save time generating walks by avoiding redundant computations for transition probabilities (**Fig. 3B**). Hence, we leave the decision for the trade off between speed and memory to the user by having both precomputation (*PreComp*) and the on-the-fly (*SparseOTF, DenseOTF*) schemes as options in the software.

### Parallelism

Finally, all the modes of our new *node2vec* implementations are fully parallelized. As mentioned earlier, the process of computing transition probabilities and the process of generating walks for each node are embarrassingly parallel. Specifically, in the process of walk generation, each node is used as a starting point for an independent random walk with fixed length (unless a dead end is reached, causing early stopping) on the network. This process is repeated multiple times depending on the input parameter specified by the user. As each of these random walks are independent, multiple walks can be performed in parallel. Similarly, the precomputation of 2nd-order transition probabilities is only dependent on the 1st-order transition probabilities. Hence, in this work, the process of walk generation and 2nd-order transition probabilities precomputation are parallelized using Numba.

### These Optimizations Lead to Better Runtime and Memory Usage

As shown in **Figures 1**, **S1**, **S2**, and **S3**, all these optimizations together provide a substantial boost in performance to PecanPy over the original implementations of *node2vec* both in terms of runtime and memory usage. This includes the OTF implementations (*SparseOTF* and *DenseOTF*) successfully handling three large networks (SSN200, GIANT-TN-c01, GIANT-TN) that the original software failed to handle. Other implementations failed to run the GIANT-TN network due to memory limitations that arise from storing 2nd-order transition probabilities. The original software failed to run SSN200 because they do not support non-integer-type node IDs. *DenseOTF* failed for SSN200 since its dense-network design requires more than 5TB of memory to create a double precision dense matrix of size 800k. However, it considerably improves memory usage and speed for large dense networks like GIANT-TN. For relatively small and sparse networks (e.g. BioGRID, BlogCatalog), using *PreComp* invariably results in faster walk generation, thus achieving an overall shorter runtime. These results underscore the importance of the three modes of PecanPy.

### Quality of Node Embeddings

Finally, to ensure the quality of node embeddings generated by our new implementations, we evaluated their use as feature vectors in node classification tasks using datasets from the original paper (BlogCatalog, PPI, Wikipedia; see *Methods*). As shown in **Figure 7**, our implementations achieve the same performance as the original Python implementation. We note that there are statistically significant differences between the area under the ROC curve (auROC) scores from the original C++ and the original Python implementations. The C++ implementation is significantly worse than Python implementation for PPI (Wilcoxon p-value = 9.786e-04) and BlogCatalog (p-value = 6.837e-03), while being significantly better for Wikipedia (p-value = 7.962e-04). These differences are likely due to the different skip-gram implementations in the C++ and Python versions.

**Figure 7.**
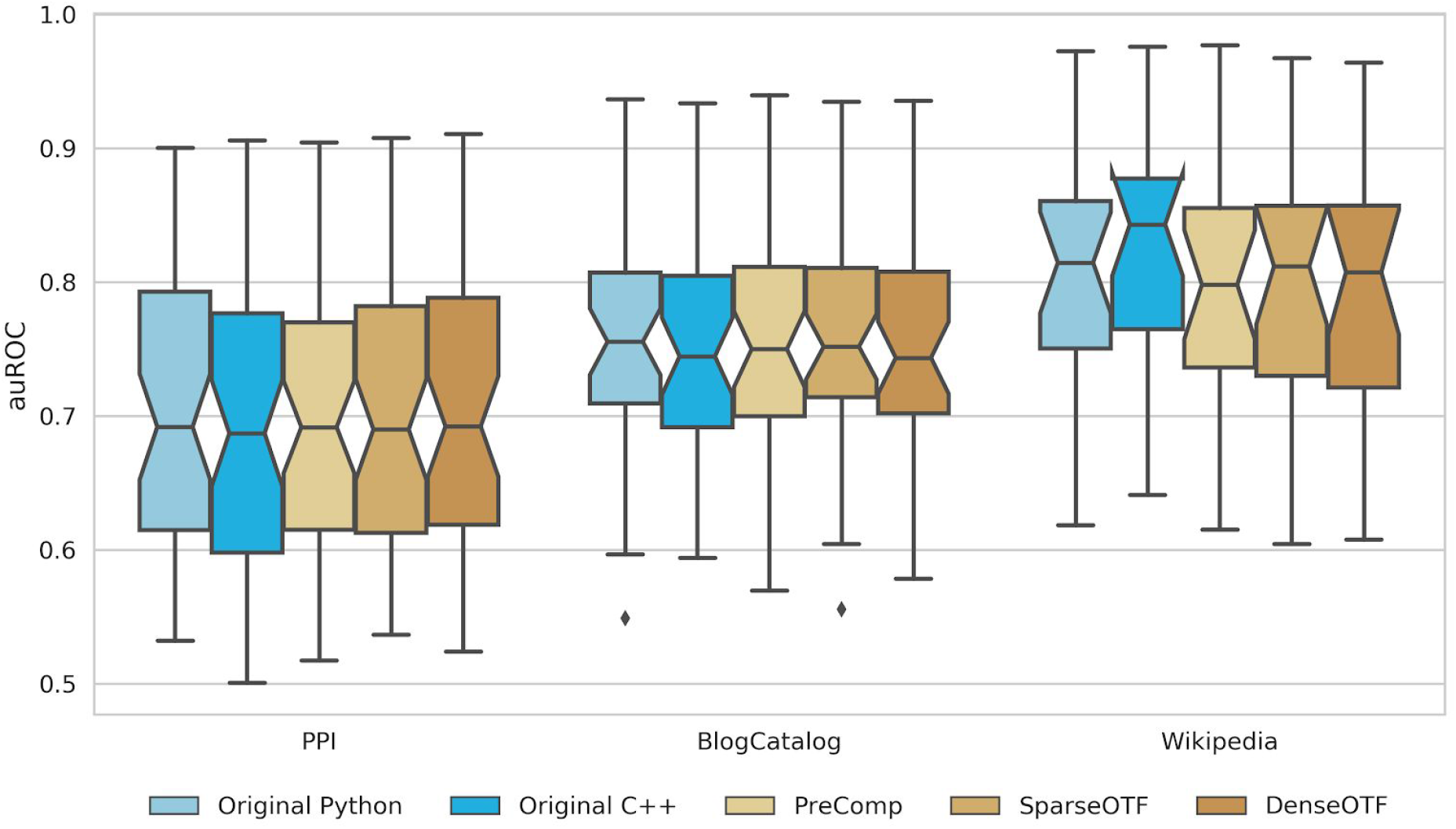
Evaluation of embedding in node classification tasks. Each group of boxplots corresponds to one of three networks. Individual boxplots in a group correspond to the distribution of auROC scores using node embeddings generated using a specific implementation (different colors).

## Discussion

Here we present PecanPy, the fastest implementation of the *node2vec* algorithm, which is the most widely used method to create numerical distributed representations of nodes in a network (graph) called node embeddings. Numerical representations of nodes open-up the possibility of deploying any tool in modern statistical or machine learning toolkit to analyze node or network properties beyond the traditional techniques of graph theory.

Node embedding is gaining rapid adoption in the analysis of biological networks (Nelson *et al.*, 2019) and *node2vec* is particularly suited for the task of node classification (Yue *et al.*, 2020).

Some examples of this general task are: 1) classifying uncharacterized genes in a functional interaction network to cellular functions they might participate in (Liu *et al.*, 2020), and 3) classifying medical terms in a term co-occurrence network (mined from electronic health records) to semantic types (e.g. drug, disease, symptoms, etc.) (Finlayson *et al.*, 2014). In all cases, with better technology and dropping costs, the ability to generate and collate massive amounts of raw data is increasing rapidly, which in turn results in ever larger networks (hundreds of thousands of nodes) with greater densities (hundreds of millions of edges) across application areas. Therefore, continuing to apply techniques such as node embedding requires software that scales reasonably with these growing large, dense networks. PecanPy serves this exact critical need.

Further, networks/graphs representations are used in nearly every discipline including sociology, economics, and language (Leskovec and Krevl, 2014). These fields are also rapidly adopting embedding-based approaches. PecanPy is a general software that can work on any network irrespective of where it comes from and what entities and relationships the nodes and edges represent. Therefore, it can find broad utility beyond biology.

## Conclusions

We have developed an efficient *node2vec* Python software – PecanPy – with significant improvement in both memory utilization and speed. We have extensively benchmarked these implementations and compared them to the original Python and C++ implementations. These analyses demonstrate that our implementations efficiently generate quality node embeddings for networks at multiple scales including large (>800k nodes) and dense (fully connected network of 26k nodes) networks that both original implementations failed to execute. The open source PecanPy software and documentation are available at https://github.com/krishnanlab/pecanpy.

## Methods

### Networks

We used a collection of eight networks for testing and benchmarking all *node2vec* implementations. PPI, BlogCatalog, and Wikipedia are from the original *node2vec* paper (Grover and Leskovec, 2016) (download link from node2vec webpage https://snap.stanford.edu/node2vec/). BioGRID (Stark *et al.*, 2006), STRING (Szklarczyk *et al.*, 2015), and GIANT-TN (Greene *et al.*, 2015) are molecular interaction networks (download from https://doi.org/10.5281/zenodo.3352323), where nodes are proteins/genes and edges are interactions between them. GIANT-TN-c01 is a sub-network of GIANT-TN where edges with edge weight below 0.01 are discarded. SSN200 (Law *et al.*, 2019) is a cross-species network of proteins from 200 species (download from https://bioinformatics.cs.vt.edu/~jeffl/supplements/2019-fastsinksource/), with the edges representing protein sequence similarities. **Table 2** contains summary statistics of the above networks including number of nodes, edges, and network density.

### Benchmarking Runtime and Memory Usage

We profiled the runtime and memory usage for each implementation on each network. All tests were performed using a 28 core Intel Xeon CPU E5-2680 v4 @2.4GHz. Each one of four stages in the *node2vec* software was timed individually. To profile the original implementations, we added timers to each stage using the built-in time function in Python and C++. Memory was profiled using the system built-in GNU timer, which could measure the maximum resident size (physical memory usage) throughout the runtime of a program. Runtime and memory profiling were carried out with two different resource configurations. The results in the main paper are based on a multi-core configuration, with 28 cores, 200GB allocated memory, and a 24-hour time limit. We also profiled the implementations in a single-core configuration, with 1 core, 32 GB memory allocated, and 8 hours time limit. The multi-core setup emulates performance on a high-performance computing facility, while the single-core setup emulates the scenario of the software being run on a personal computer. All testing results can be found in **Table S1**.

### Evaluating the Quality of Node Embeddings

The three networks used in the *node2vec* paper (PPI, BlogCatalog, Wikipedia) come with labels associated with each node. Performance of node classification tasks using the generated embeddings from different implementations are compared using the three networks. Some of the labelsets from the data repository appeared to have very few positive examples (e.g. 1). For more rigorous evaluation, only labelsets with at least 10 positive examples are evaluated. For each labelset, a one vs rest L2 regularized logistic regression model is trained on the training set and tested on the testing set, with 5-fold cross validation, repeated 10 times. Area under receiver operating curves (auROC) is used as the performance metrics. For each labelset, the median value of auROC among 10 repeated 5-fold cross validation is reported as the final score for each labelset-implementation pair. For each network, and for each implementation, a wilcoxon paired test is performed against the original Python implementation.

## Declarations

### Ethics approval and consent to participate

Not applicable

### Consent for publication

Not applicable

### Availability of data and materials

The networks used in the original node2vec publication are here: https://snap.stanford.edu/node2vec/. The SSN200 network is available at https://bioinformatics.cs.vt.edu/~jeffl/supplements/2019-fastsinksource/. All other networks are available at https://doi.org/10.5281/zenodo.3352323. The PecanPy software, along with extensive documentation on how to run it, is available at https://github.com/krishnanlab/pecanpy.

### Authors’ contributions

RL and AK designed the study. RL wrote the software and performed all the analyses. RL and AK interpreted the results and wrote the final manuscript.

## Acknowledgements

We thank Christopher A. Mancuso, Anna Yannakopoulos, and the rest of the Krishnan Lab for valuable discussions and feedback on the software and manuscript.

