## Supplementary Materials for "PecanPy: a fast, efficient, and parallelized Python implementation of *node2vec*"

### 1. Supplementary Tables

**Table S1. All Runtimes for All Implementations on All Networks**. The entries for Total time and Maximum residence size that are colored dark grey indicate cases when the implementation in the row failed, at which point these data (time and size) were recorded.

| Network | Method | Setup | Loading time | Preproce -ssing time | Walking time | Training time | Total time | Total time in second | Maximum residence size |
|---|---|---|---|---|---|---|---|---|---|
| BioGRID | Original Python | Multi | 0:3.3 | 1:44.70 | 8:29.85 | 0:24.12 | 10:47.78 | 647.78 | 2.6GB |
| BioGRID | Original C++ | Multi | 0:1 | 0:1 | 0:1 | 2:40.00 | 2:45.03 | 165.03 | 1.0GB |
| BioGRID | PecanPy - PreComp | Multi | 0:0.64 | 0:5.71 | 0:7 | 0:22.36 | 0:37.21 | 37.21 | 1.2GB |
| BioGRID | PecanPy - SparseOTF | Multi | 0:0.61 | 0:0 | 0:52.48 | 0:24.45 | 1:19.04 | 79.04 | 407.3MB |
| BioGRID | PecanPy - DenseOTF | Multi | 0:5.13 | 0:0 | 1:09.44 | 0:24.47 | 1:41.03 | 101.03 | 3.9GB |
| BioGRID | Original Python | Single | 0:3.8 | 2:00.37 | 9:47.81 | 3:30.41 | 15:39.97 | 939.97 | 2.6GB |
| BioGRID | Original C++ | Single | 0:1 | 0:15 | 0:7 | 5:49.00 | 6:13.61 | 373.61 | 1.0GB |
| BioGRID | PecanPy - PreComp | Single | 0:0.77 | 0:37.51 | 0:10.51 | 4:22.56 | 5:15.19 | 315.19 | 1.2GB |
| BioGRID | PecanPy - SparseOTF | Single | 0:0.89 | 0:0 | 0:55.17 | 4:14.52 | 5:12.96 | 312.96 | 376.1MB |
| BioGRID | PecanPy - DenseOTF | Single | 0:6.63 | 0:0 | 24:54.10 | 4:22.65 | 29:26.23 | 1766.23 | 3.9GB |
| BlogCatalog | Original Python | Multi | 0:4.12 | 8:14.21 | 10:47.90 | 0:12.86 | 19:25.52 | 1165.52 | 6.5GB |
| BlogCatalog | Original C++ | Multi | 0:1 | 0:8 | 0:0 | 1:17.00 | 1:27.98 | 87.98 | 4.3GB |
| BlogCatalog | PecanPy - PreComp | Multi | 0:0.73 | 0:6.28 | 0:4.64 | 0:11.99 | 0:25.02 | 25.02 | 4.4GB |
| BlogCatalog | PecanPy - SparseOTF | Multi | 0:0.7 | 0:0 | 0:28.55 | 0:12.08 | 0:42.71 | 42.71 | 293.5MB |
| BlogCatalog | PecanPy - DenseOTF | Multi | 0:1.4 | 0:0 | 0:29.1 | 0:11.98 | 0:44.42 | 44.42 | 1.2GB |
| BlogCatalog | Original Python | Single | 0:4.85 | 9:31.43 | 12:10.09 | 1:41.30 | 23:46.38 | 1426.38 | 6.5GB |
| BlogCatalog | Original C++ | Single | 0:1 | 1:09.00 | 0:4 | 2:44.00 | 3:59.74 | 239.74 | 4.3GB |
| BlogCatalog | PecanPy - PreComp | Single | 0:0.94 | 0:40.17 | 0:6.7 | 1:54.43 | 2:46.01 | 166.01 | 4.4GB |
| BlogCatalog | PecanPy - SparseOTF | Single | 0:0.99 | 0:0 | 0:53.66 | 1:54.20 | 2:51.16 | 171.16 | 269.7MB |

| | | | | | | | | | |
|---|---|---|---|---|---|---|---|---|---|
| BlogCatalog | PecanPy - DenseOTF | Single | 0:1.87 | 0:0 | 8:30.50 | 1:55.28 | 10:30.40 | 630.40 | 1.1GB |
| GIANT-TN-c01 | Original Python | Multi | - | - | - | - | 10:12:59 | 36779.00 | 200.0GB |
| GIANT-TN-c01 | Original C++ | Multi | - | - | - | - | 12:16.57 | 736.57 | 200.0GB |
| GIANT-TN-c01 | PecanPy - PreComp | Multi | - | - | - | - | 2:50.13 | 170.13 | 4.8GB |
| GIANT-TN-c01 | PecanPy - SparseOTF | Multi | 1:34.82 | 0:0 | 1:12.47 | 0:25.81 | 3:14.75 | 194.75 | 4.8GB |
| GIANT-TN-c01 | PecanPy - DenseOTF | Multi | 8.28 | 0:0 | 3:12.19 | 0:27 | 3:55.76 | 235.76 | 6.0GB |
| GIANT-TN-c01 | Original Python | Single | - | - | - | - | 1:49:32 | 6572.00 | 32.0GB |
| GIANT-TN-c01 | Original C++ | Single | - | - | - | - | 9:38.43 | 578.43 | 32.0GB |
| GIANT-TN-c01 | PecanPy - PreComp | Single | - | - | - | - | 3:43.21 | 223.21 | 4.8GB |
| GIANT-TN-c01 | PecanPy - SparseOTF | Single | 2:08.15 | 0:0 | 10:52.75 | 5:29.50 | 18:32.87 | 1112.87 | 4.8GB |
| GIANT-TN-c01 | PecanPy - DenseOTF | Single | 0:10.36 | 0:0 | 1:20:13 | 5:33.62 | 1:26:00 | 5160.00 | 5.9GB |
| GIANT-TN | Original Python | Multi | - | - | - | - | - | - | - |
| GIANT-TN | Original C++ | Multi | - | - | - | - | 6:14:36 | 22476.00 | 200.0GB |
| GIANT-TN | PecanPy - PreComp | Multi | - | - | - | - | 20:32.58 | 1232.58 | 39.3GB |
| GIANT-TN | PecanPy - SparseOTF | Multi | 10:27.14 | 0:0 | 2:53.77 | 0:27.3 | 13:50.62 | 830.62 | 39.3GB |
| GIANT-TN | PecanPy - DenseOTF | Multi | 0:8.57 | 0:0 | 2:48.07 | 0:27.35 | 3:27.70 | 207.70 | 6.0GB |
| GIANT-TN | Original Python | Single | - | - | - | - | 10:09.78 | 609.78 | 32.0GB |
| GIANT-TN | Original C++ | Single | - | - | - | - | 6:35:01 | 23701.00 | 32.0GB |
| GIANT-TN | PecanPy - PreComp | Single | - | - | - | - | 10:46.71 | 646.71 | 32.0GB |
| GIANT-TN | PecanPy - SparseOTF | Single | - | - | - | - | 11:00.76 | 660.76 | 32.0GB |
| GIANT-TN | PecanPy - DenseOTF | Single | 0:10.43 | 0:0 | 1:09:56 | 5:48.38 | 1:15:58 | 4558.00 | 6.0GB |
| PPI | Original Python | Multi | 0:0.52 | 0:8.6 | 0:45.94 | 0:4.18 | 1:03.37 | 63.37 | 423.4MB |
| PPI | Original C++ | Multi | 0 | 0:0 | 0:0 | 0:31 | 0:31.79 | 31.79 | 110.9MB |
| PPI | PecanPy - PreComp | Multi | 0:0.11 | 0:3.78 | 0:3.62 | 0:4.85 | 0:14.02 | 14.02 | 317.0MB |

| | | | | | | | | | |
|---|---|---|---|---|---|---|---|---|---|
| PPI | PecanPy - SparseOTF | Multi | 0:0.1 | 0:0 | 0:13.35 | 0:4.62 | 0:19.42 | 19.42 | 252.6MB |
| PPI | PecanPy - DenseOTF | Multi | 0:0.26 | 0:0 | 0:12.56 | 0:4.97 | 0:19.15 | 19.15 | 338.6MB |
| PPI | Original Python | Single | 0:0.59 | 0:10.55 | 0:56.05 | 0:34.04 | 1:57.67 | 117.67 | 418.5MB |
| PPI | Original C++ | Single | 0:0 | 0:1 | 0:0 | 0:59 | 1:01.23 | 61.23 | 109.8MB |
| PPI | PecanPy - PreComp | Single | 0:0.13 | 0:9.78 | 0:4.65 | 0:38.62 | 0:56.88 | 56.88 | 293.8MB |
| PPI | PecanPy - SparseOTF | Single | 0:0.11 | 0:0 | 0:11.1 | 0:31.78 | 0:45.34 | 45.34 | 224.2MB |
| PPI | PecanPy - DenseOTF | Single | 0:0.34 | 0:0 | 1:03.94 | 0:39.25 | 1:46.24 | 106.24 | 318.0MB |
| SSN200 | Original Python | Multi | - | - | - | - | 0:3.23 | 3.23 | 69.5MB |
| SSN200 | Original C++ | Multi | - | - | - | - | 0:0.05 | 0.05 | 1.9MB |
| SSN200 | PecanPy - PreComp | Multi | - | - | - | - | 1:24:50 | 5090.00 | 182.2GB |
| SSN200 | PecanPy - SparseOTF | Multi | 3:24.12 | 0:0 | 33:48.12 | 18:09.47 | 55:31.35 | 3331.35 | 10.7GB |
| SSN200 | PecanPy - DenseOTF | Multi | - | - | - | - | 0:1.28 | 1.28 | 109.2MB |
| SSN200 | Original Python | Single | - | - | - | - | 0:16.31 | 16.31 | 69.5MB |
| SSN200 | Original C++ | Single | - | - | - | - | 0:0.2 | 0.20 | 1.9MB |
| SSN200 | PecanPy - PreComp | Single | - | - | - | - | 3:45.99 | 225.99 | 32.0GB |
| SSN200 | PecanPy - SparseOTF | Single | 4:00.10 | 0:0 | 39:26.41 | 3:12:39 | 3:56:11 | 14171.00 | 10.7GB |
| SSN200 | PecanPy - DenseOTF | Single | - | - | - | - | 0:2.57 | 2.57 | 109.4MB |
| STRING | Original Python | Multi | 1:00.22 | 4:44:11 | 45:34.18 | 0:21.53 | 5:31:41 | 19901.00 | 112.1GB |
| STRING | Original C++ | Multi | 0:6 | 3:20.00 | 0:2 | 1:48.00 | 5:44.24 | 344.24 | 80.9GB |
| STRING | PecanPy - PreComp | Multi | 0:8.38 | 0:37.29 | 0:6.31 | 0:18.02 | 1:11.74 | 71.74 | 80.1GB |
| STRING | PecanPy - SparseOTF | Multi | 0:8.47 | 0:0 | 0:46.86 | 0:18.12 | 1:14.98 | 74.98 | 560.3MB |
| STRING | PecanPy - DenseOTF | Multi | 0:3.8 | 0:0 | 1:03.82 | 0:19.33 | 1:28.42 | 88.42 | 2.9GB |
| STRING | Original Python | Single | - | - | - | - | 2:04:38 | 7478.00 | 32.0GB |
| STRING | Original C++ | Single | - | - | - | - | 0:48.89 | 48.89 | 32.0GB |

| | | | | | | | | | |
|---|---|---|---|---|---|---|---|---|---|
| STRING | PecanPy - PreComp | Single | - | - | - | - | 0:29.84 | 29.84 | 32.0GB |
| STRING | PecanPy - SparseOTF | Single | 0:10.44 | 0:0 | 2:13.42 | 3:39.82 | 6:06.03 | 366.03 | 561.0MB |
| STRING | PecanPy - DenseOTF | Single | 0:4.17 | 0:0 | 20:52.40 | 2:43.73 | 23:43.21 | 1423.21 | 2.9GB |
| Wikipedia | Original Python | Multi | 0:2.33 | 3:27.54 | 4:31.34 | 0:5.89 | 8:12.34 | 492.34 | 1.9GB |
| Wikipedia | Original C++ | Multi | 0:0 | 0:2 | 0:0 | 0:39 | 0:41.73 | 41.73 | 1.2GB |
| Wikipedia | PecanPy - PreComp | Multi | 0:0.36 | 0:4.2 | 0:3.98 | 0:6.22 | 0:16.06 | 16.06 | 1.4GB |
| Wikipedia | PecanPy - SparseOTF | Multi | 0:0.33 | 0:0 | 0:15.56 | 0:5.52 | 0:22.75 | 22.75 | 262.3MB |
| Wikipedia | PecanPy - DenseOTF | Multi | 0:0.39 | 0:0 | 0:14.38 | 0:5.71 | 0:22.19 | 22.19 | 419.7MB |
| Wikipedia | Original Python | Single | 0:2.79 | 4:34.85 | 5:31.11 | 0:30.73 | 10:56.31 | 656.31 | 1.9GB |
| Wikipedia | Original C++ | Single | 0:1 | 0:18 | 0:2 | 1:13.00 | 1:34.02 | 94.02 | 1.2GB |
| Wikipedia | PecanPy - PreComp | Single | 0:0.43 | 0:14.85 | 50:0.46 | 0:33.62 | 0:56.6 | 56.60 | 1.3GB |
| Wikipedia | PecanPy - SparseOTF | Single | 0:0.44 | 0:0 | 0:42.4 | 0:33.93 | 1:18.98 | 78.98 | 230.1MB |
| Wikipedia | PecanPy - DenseOTF | Single | 0:0.55 | 0:0 | 2:35.32 | 0:33.05 | 3:12.61 | 192.61 | 394.8MB |

### 2. Supplementary Figures

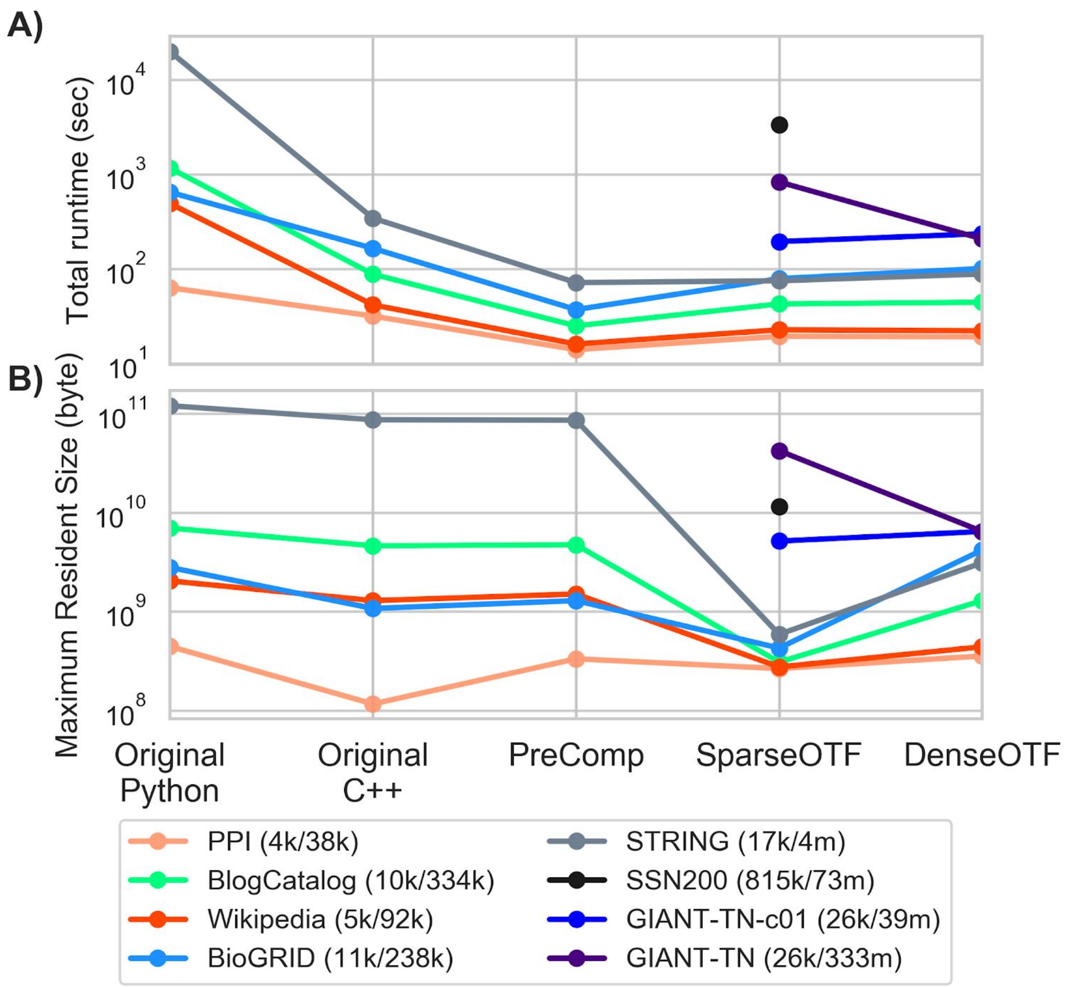


**Figure S1. Performance of the original Python and C++ implementations and the three new implementations –** ***PreComp*****,** ***SparseOTF*****, and** ***DenseOTF*** **– on eight networks using multiple cores.** The parallel plots trace the performance of different *node2vec* implementations (x-axis) for 8 networks (colored dots/lines; number of nodes/edges are in legend below) in terms of (A) total runtime (seconds) and (B) peak memory usage (bytes). Absence of a dot indicates the failure of a particular implementation to run for a particular network.

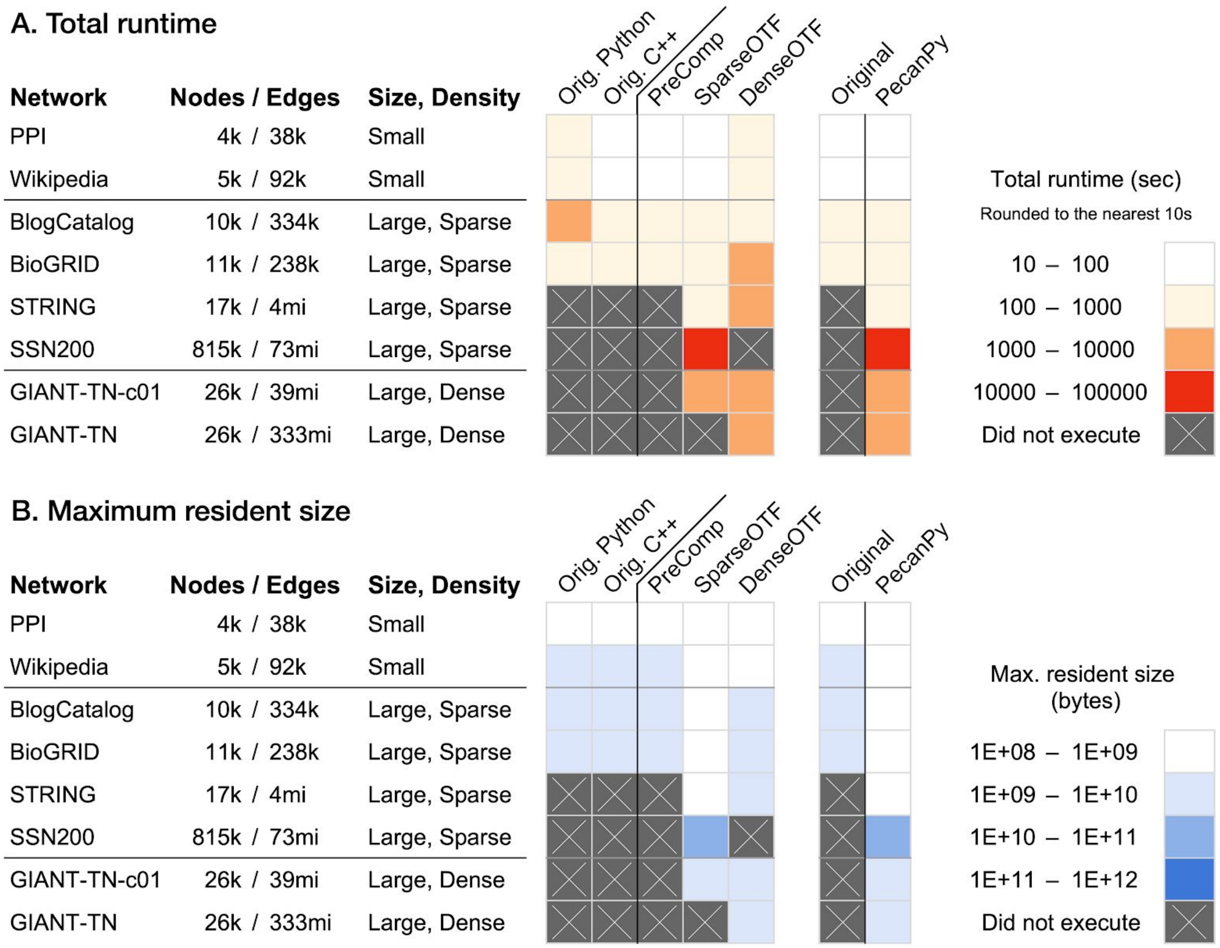


**Figure S2. Summary of runtime and memory of PecanPy and the original implementations of *node2vec* using a single core.** The eight networks of varying sizes and densities are along the rows. The software implementations are along the columns. The first heatmap (on the left) shows the performance of the original Python and C++ software along with the three modes of PecanPy (*PreComp*, *SparseOTF*, and *DenseOTF*). The adjacent 2-column heatmap (on the right) summarizes the performance of the original (best of Python and C++ versions) and PecanPy (best of *PreComp*, *SparseOTF*, and *DenseOTF*) implementations. Lighter colors correspond to lower runtime in panel A and lower memory usage in panel B. Crossed grey indicates that the particular implementation (column) failed to run for a particular network (row).

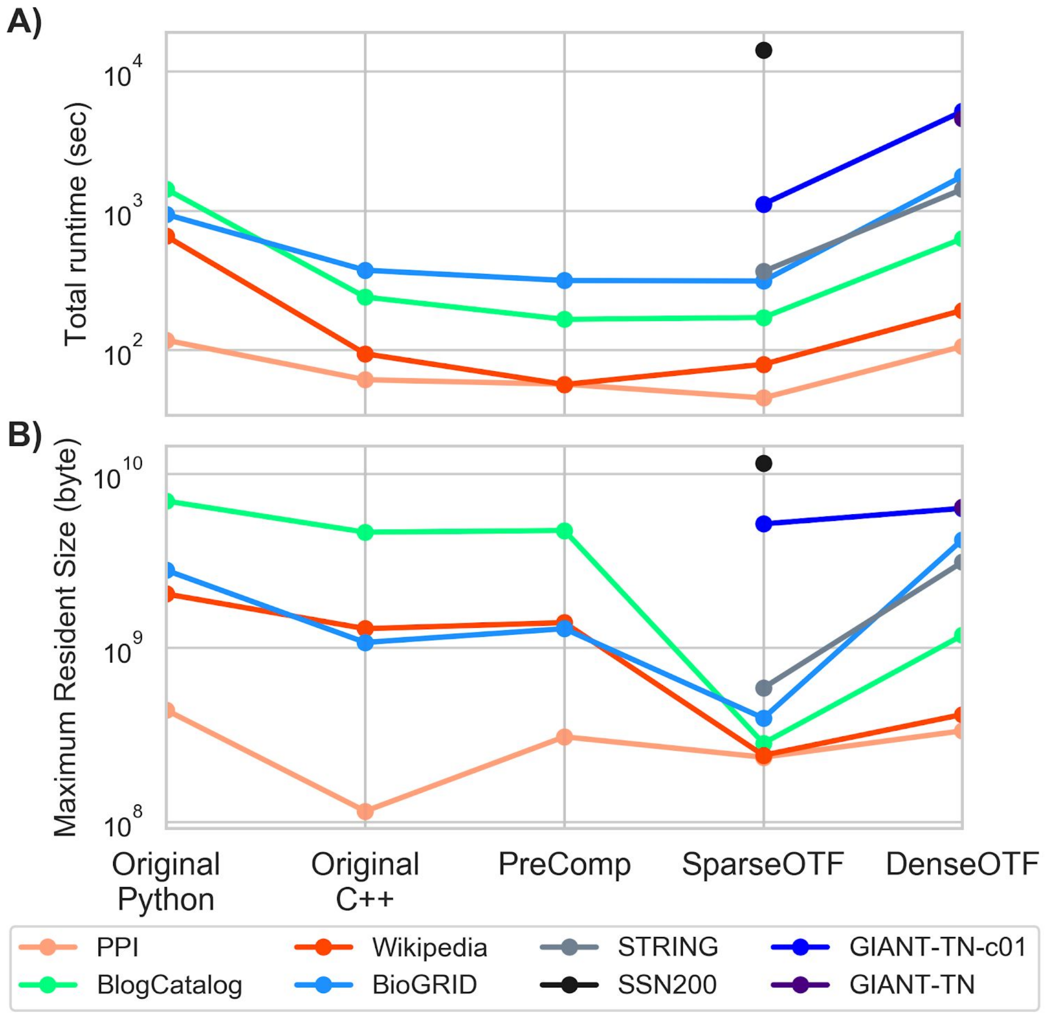


**Figure S3. Performance of the original Python and C++ implementations and the three new implementations – *PreComp*, *SparseOTF*, and *DenseOTF* – on eight networks using a single core.** The parallel plots trace the performance of different *node2vec* implementations (x-axis) for 8 networks (colored dots/lines) in terms of (A) total runtime (seconds) and (B) peak memory usage (bytes). Absence of a dot indicates the failure of a particular implementation to run for a particular network.

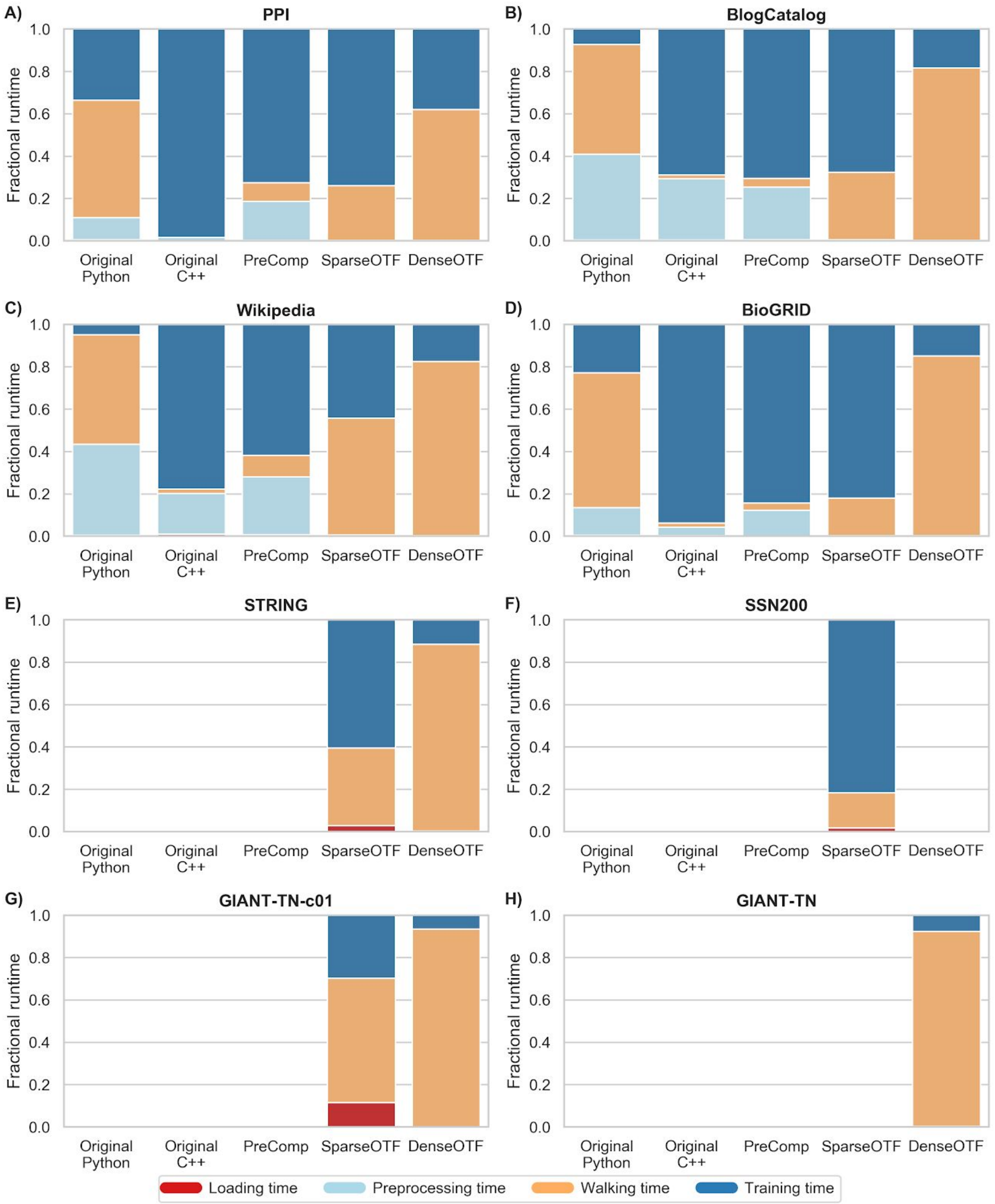


**Figure S4. Fraction of runtime contributed by each stage of *node2vec* in different implementations using a single core.** Each panel corresponds to a single network and each stacked bar within a panel corresponds to an individual *node2vec* implementation. The height of each segment within a bar represents the fraction of runtime contributed by each of the different stages of *node2vec*, tested in a single-core configuration.

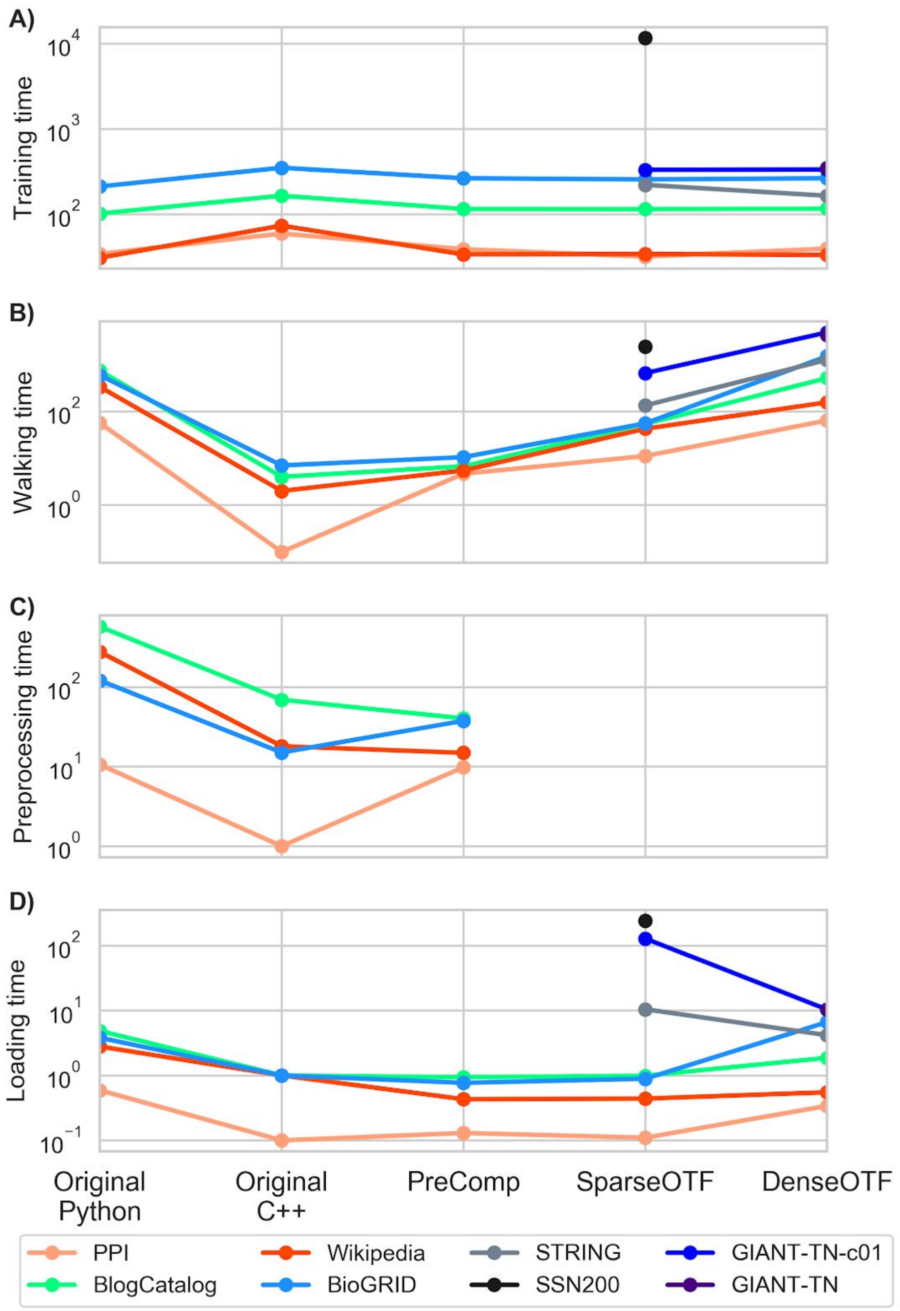


**Figure S5. Raw runtimes of each stage of *node2vec* in different implementations using a single core.** Each parallel plot corresponds to one of four stages of *node2vec*. Each line traces the raw runtime (points on parallel y-axes) of a specific network (color) across the different implementations (x-axis). In all plots, absence of a point for a particular network for any implementation indicates that the network failed to load.
